# Helical Coiled Nucleosome Chromosome Architectures during Cell Cycle Progression

**DOI:** 10.1101/2024.05.25.595892

**Authors:** Angus McDonald, Cornelis Murre, John Sedat

**Affiliations:** Micron School of Materials Science and Engineering, Boise State University, Boise, ID 83725-2090; Division of Biological Sciences, University of California San Diego, La Jolla, CA 92093; Department of Biochemistry and Biophysics, University of California, San Francisco, San Francisco, CA 94158

**Keywords:** Nuclear Structure, Chromosome Architecture, Computer Modeling, and Cell Cycle

## Abstract

Recent studies showed an interphase chromosome architecture, --- a specific coiled nucleosome structure, --- derived from cryo-preserved EM tomograms, and dispersed throughout the nucleus. The images were computationally processed to fill in the missing wedges of data caused by incomplete tomographic tilts. The resulting structures increased z-resolution enabling an extension of the proposed architecture to that of mitotic chromosomes.

Here we provide additional insights and details into the coiled nucleosome chromosome architectures. We build on the defined chromosomes time-dependent structures in an effort to probe their dynamics. Variants of the coiled chromosome structures, possibly further defining specific regions, are discussed. We propose, based on generalized specific uncoiling of mitotic chromosomes in telophase, large-scale re-organization of interphase chromosomes. Chromosome territories, organized as micron-sized small patches, are constructed, satisfying complex volume considerations. Finally, we unveiled the structures of replicated coiled chromosomes, still attached to centromeres, as part of chromosome architecture.

**Significance Statement:** This study places all 46 sequenced human chromosomes, --- correctly filled with nucleosomes and in micron sized chromosome territories — into 10micron (average sized) nuclei. The chromosome architecture used a helical nucleosome coiled structure discerned from cryo-EM tomography, as was recently published (https://doi.org/10.1073/pnas.2119101119). This chromosome architecture was further modeled to dynamic structures, structure variations and chromosome replication centromere complications. Finally, this chromosome architecture was modified to allow seamless transition through the cell cycle.

## Introduction and Results

The dominant current paradigm for interphase chromosome structure within the nucleus is a flexible polymer-based system, following polymer statistics (1-6 & refs therein). Nevertheless, there is a degree of order as high throughput DNA sequencing of chromosome conformation captured chromatin have shown (7-9 & refs therein). In addition, the relatively specific site associations can be linked to genetic function and control, including during cell type specification and development (10-12).

An alternative depiction of the interphase chromosome structure can be derived from cryo-EM studies. The cell with its nucleus is rapidly frozen, at liquid nitrogen temperatures, to a glassy ice, preserving the interior proteins, nucleic acids, membranes, and water (13-17). The EM data is computer processed, using deconvolution methodology to fill-in the missing high angle data coming from the incomplete tilts (18,19). The increased Z resolution together with 3-Dimensional visualization, as stereo rocking movies, now allows insights into the preserved dense nuclear interior ( https://doi.org/10.1073/pnas.2119101119,20). These studies indicated that interphase chromosomes fold into an ∼200nm diameter helically coiled nucleosome fiber, termed a Slinky (defined as interphase S --20,21). Euchromatin-transcribed sequences --appear to be extended interphase S, while heterochromatin folds into a more compact interphase S structure (20). Subsequent studies showed an architecture for prophase and mitotic chromosomes, as modifications of additional coiling processes (https://doi.org/10.1073/pnas.2119107119,22). Prophase structure is now termed --prophase S’, with mitotic chromosome architecture as -- mitotic S’’.

Fundamental to interphase chromosome structure as well as prophase and mitotic chromosomes are hard boundary conditions ---The known facts. We examined human chromosomes as a model system. All 46 chromosomes are fully DNA sequenced (23). Since approximately 90% of the DNA is structured as nucleosomes (24,25), with one nucleosome per 200 base pairs, likely organized as an 11nm nucleosome fiber, each chromosome will have an essentially defined nucleosome total (Table 1). The essentially complete filling of a given chromosome DNA length with nucleosomes as described above has to be qualified. This result comes from analysis of nucleosome packing in rat liver nuclei. It is possible that other specific cell types will have less nucleosome density (23,25). The interphase chromosomes have to fit, as micron-sized chromosome territories, into a nucleus of about 10microns diameter. Likewise, mitotic chromosomes have dimensions observed from mitotic chromosome spreads (26-29). These boundary limits will restrict what architectures are possible, as were used in the proposed human mitotic chromosome 10 structure (22).

**Table 1.**
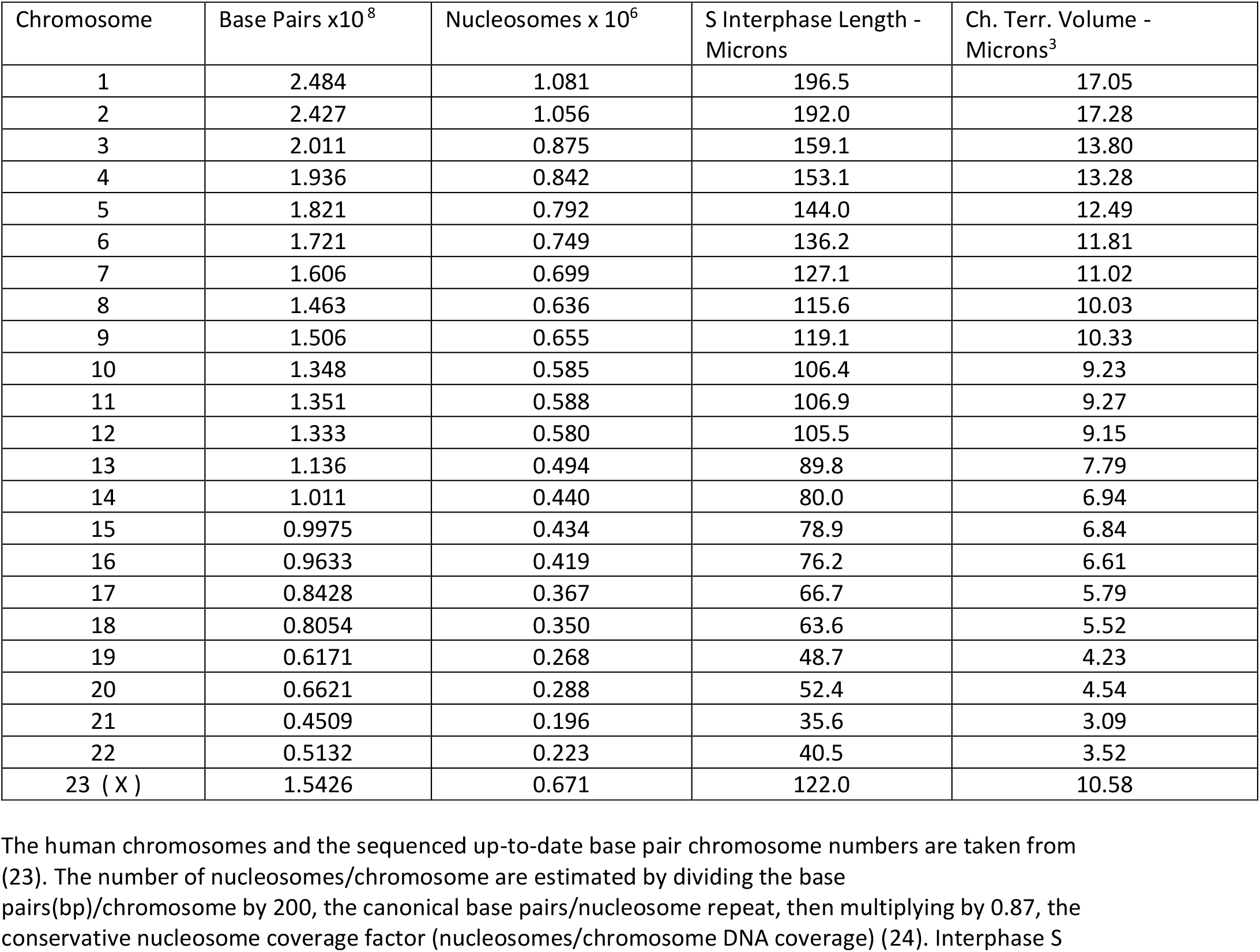
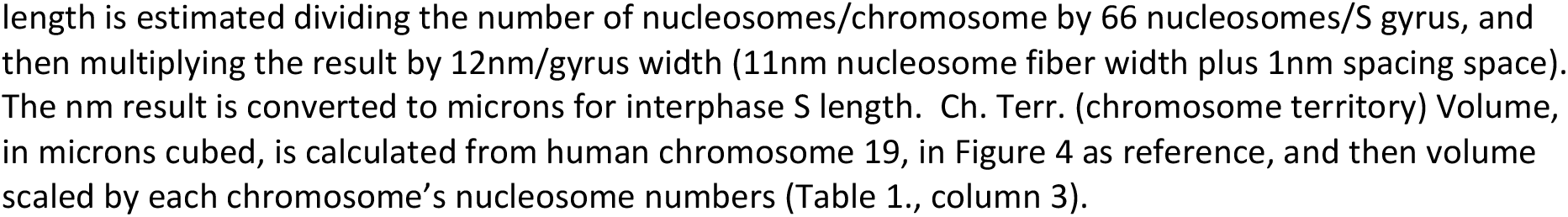
The numbered human chromosomes, sequenced length in base pairs, number of nucleosomes / length, the S interphase length, and Ch. Terr. Vol. are shown in Table 1.

| Chromosome | Base Pairs x10<sup>8</sup> | Nucleosomes x 10<sup>6</sup> | S Interphase Length - Microns | Ch. Terr. Volume - Microns<sup>3</sup> |
| --- | --- | --- | --- | --- |
| 1 | 2.484 | 1.081 | 196.5 | 17.05 |
| 2 | 2.427 | 1.056 | 192.0 | 17.28 |
| 3 | 2.011 | 0.875 | 159.1 | 13.80 |
| 4 | 1.936 | 0.842 | 153.1 | 13.28 |
| 5 | 1.821 | 0.792 | 144.0 | 12.49 |
| 6 | 1.721 | 0.749 | 136.2 | 11.81 |
| 7 | 1.606 | 0.699 | 127.1 | 11.02 |
| 8 | 1.463 | 0.636 | 115.6 | 10.03 |
| 9 | 1.506 | 0.655 | 119.1 | 10.33 |
| 10 | 1.348 | 0.585 | 106.4 | 9.23 |
| 11 | 1.351 | 0.588 | 106.9 | 9.27 |
| 12 | 1.333 | 0.580 | 105.5 | 9.15 |
| 13 | 1.136 | 0.494 | 89.8 | 7.79 |
| 14 | 1.011 | 0.440 | 80.0 | 6.94 |
| 15 | 0.9975 | 0.434 | 78.9 | 6.84 |
| 16 | 0.9633 | 0.419 | 76.2 | 6.61 |
| 17 | 0.8428 | 0.367 | 66.7 | 5.79 |
| 18 | 0.8054 | 0.350 | 63.6 | 5.52 |
| 19 | 0.6171 | 0.268 | 48.7 | 4.23 |
| 20 | 0.6621 | 0.288 | 52.4 | 4.54 |
| 21 | 0.4509 | 0.196 | 35.6 | 3.09 |
| 22 | 0.5132 | 0.223 | 40.5 | 3.52 |
| 23 ( X ) | 1.5426 | 0.671 | 122.0 | 10.58 |

The human chromosomes and the sequenced up-to-date base pair chromosome numbers are taken from (23). The number of nucleosomes/chromosome are estimated by dividing the base pairs(bp)/chromosome by 200, the canonical base pairs/nucleosome repeat, then multiplying by 0.87, the conservative nucleosome coverage factor (nucleosomes/chromosome DNA coverage) (24). Interphase S

### Recent Kinetic Extensions of Interphase S, Prophase S’ and Mitotic S’’ Chromosome Structures

We extend the analysis of the coiled chromosome structures described in our recent studies (20,22). The Slinky structures, interphase S, prophase S’ and mitotic S’’, are now built as a time series showing these structures coiling (Figure 1). We use an engineering coiling software (30), suitable modified, so that the correctly dimensioned molecularly accurate structures are shown and analyzed as in previous studies (20,22). This time series allows one to search for difficult building and/or formation points, including kinks, compression points, or abrupt bends in dynamic coiling. Study of the kinetic data, suggests a smooth construction series without obvious problems (Figure. 1). By monitoring the kinetics displayed by movies in a backward manner, we were able to visualize interphase S structure pulling apart, to different degrees, during the course of nascent transcription.

**Figure 1.**
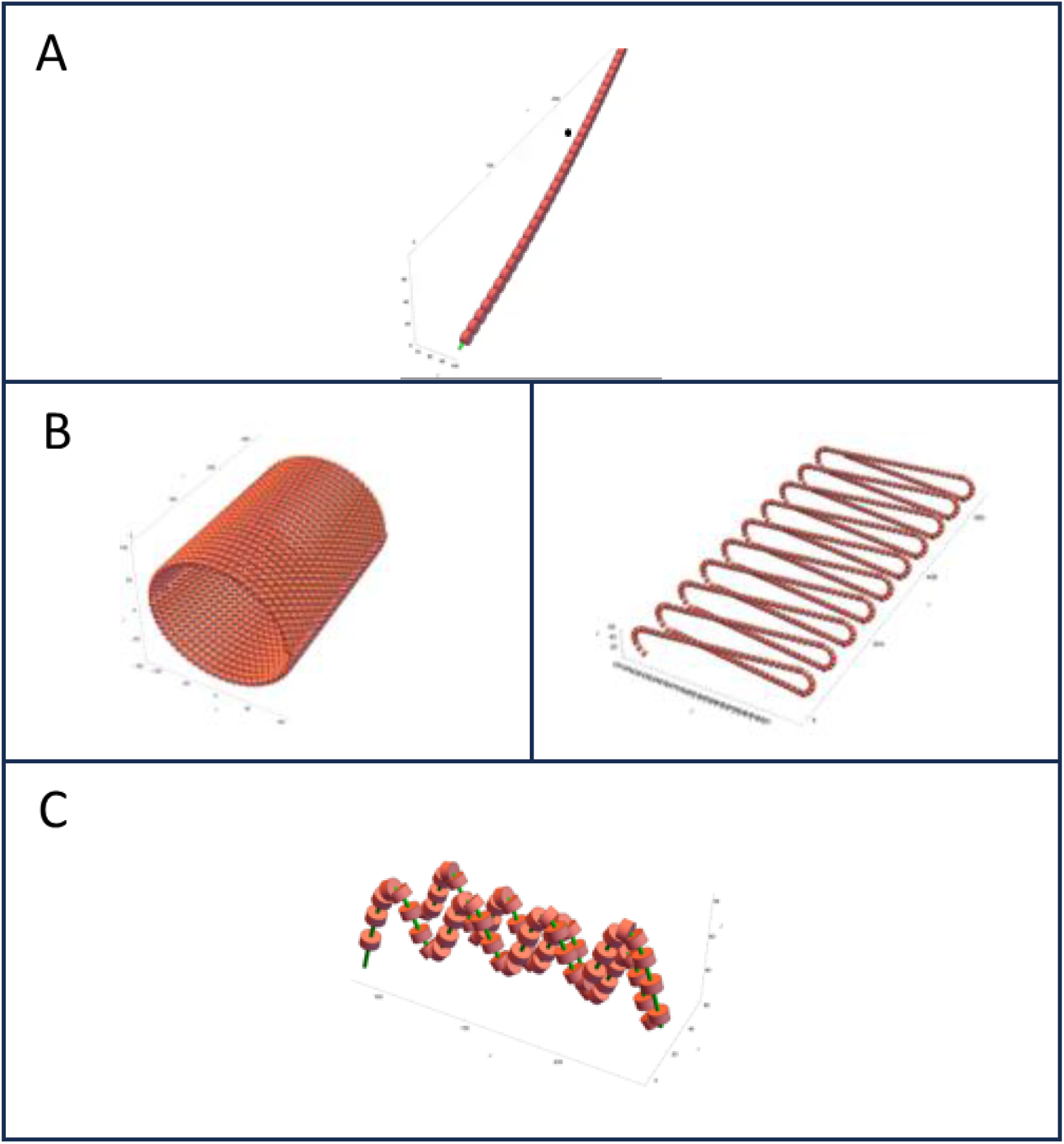
A unified chromosome architecture based on multiple helical structure coiling as a dynamic process. Panel A shows the helical coiling of the 11nm nucleosome fiber into the interphase chromosome, termed interphase S, a structure extensively described in (20). This figure emphasizes the coiling dynamics as clickable/link movies—activated by clicking on the structure, and the coiling controlled by the movie bar(red) left-to-right for coiling and right-to-left for uncoiling. [note the right-hand side of the movie bar controls sound, not used] The more open coiling, initially, and later the pulled-out coils emphasize the transcription process, while the more compressed coiling is likely heterochromatin structure, best seen in Figure 1B. Note that enlarged views of the movies are possible by clicking the enlargement button. Some practice, playing the movies, shows additional features of the dynamics.

One of the main reasons for developing the kinetic series is a proposal to use the time series to intersect with Finite Element Analysis (FEA) (31,32), the detailed modern physics quantitation and analysis approach. With this methodology, it is possible to determine fundamental forces, and estimate diffusion parameters in solvents of suitable viscosity. Could bending, coiling forces, and forces required to pull the interphase S gyri apart for transcription be determined? What are the forces to indent and distort the prophase S and mitotic S’ gyri? What forces are required to coil the mitotic S’’ structure in the final mitotic helical winding? Could handedness preference, at the different levels of coiling, be investigated? The FEA approach should yield physics-based data that can be linked to biochemical study for the investigation of nuclear and chromosome structure. Experimental data from optical and Cryo-EM tomography should intersect with the FEA approach, and is essential to achieve a deeper understanding of chromosome structure.

It should be noted that there are several ways to coil structures; the approach described above emphasizes search for coiling problems and a possible link to the proposed FEA direction. Another coiling paradigm, coiling an entire intact built structure, with time points, is described in Figure 3.

### Structure Variations for Interphase, Prophase and Mitotic Chromosomes

Structural variations, are seen throughout interphase S, prophase S’ and mitotic S’’ chromosomes (Figure 2). We found that the diameter of the interphase S fiber fluctuates around an average value of 200 nm but spanning distances of 100-300 nm (20) (Figure. 2). In addition, we detected many instances of indentations of the diameter of the helix (Figure 2D). A prominent example of such indentations and variations are shown in a drawing of an interphase chromosome revealing the many surface variations resulting from the interphase S diameter distinctions (Figure. 2E). We predict the presence of regions of interphase S regions that are depleted for nucleosomes that span regulatory elements including promoters and enhancer regions (33-37), and thus these depleted regions could account for some variations. Prophase S’ dimension variations, where the X axis can be extended slightly, broadened, by modulation of the indentations of the flattened prophase S’ gyri are readily revealed (Figure 2F, G). The prophase S’ width variants are further coiled to assemble mitotic chromosome resulting in mitotic S’’ helical coiling (Figure 2 H & I.). Mitotic S helical coiling resulted in an increase in the outside diameter of the mitotic coil but did not reveal changes in the inside diameter dimensions (Figure 2 H). We note that the consequence of this variation is the appearance of outside mitotic chromosome surfaces, in chromosome spreads, as bumpy/rough—many small ridges arranged roughly at 90 degrees to the chromosome long axis--because of the diameter variations. A possible rationale for the structure variations is analyzed in the Discussion.

**Figure 2.**
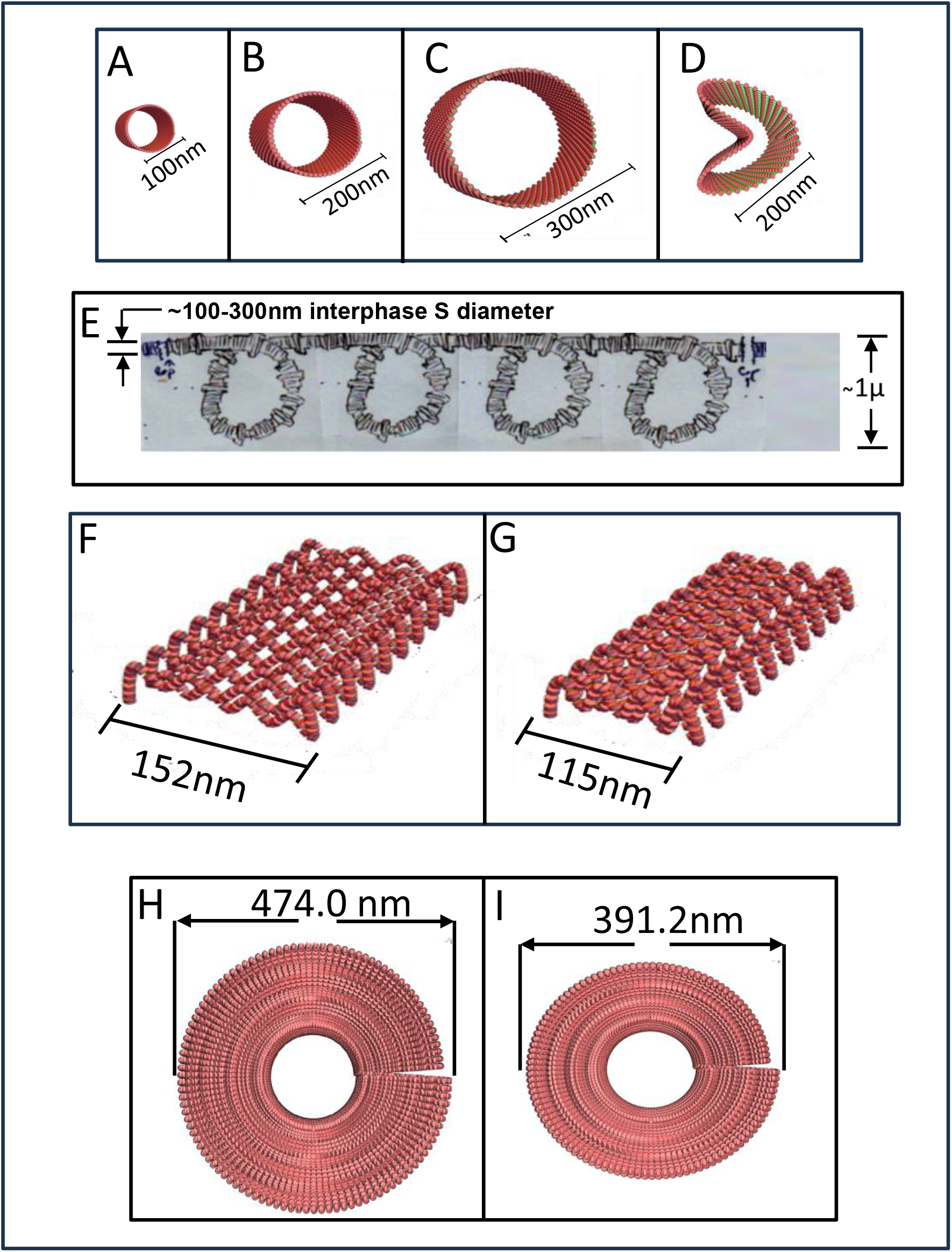
Structure variations are seen at every level of the helical coiling architecture. Figure 2. shows variation of the interphase helix (interphase S) diameter seen and documented in (20). Figure 2A. shows a 100nm diameter, the middle, Figure 2B, the most common 200nm diameter, followed by examples of 300nm diameter dimensions (Figure 2C.) The right side (Figure 2D) shows that the helix diameters were frequently misshapen and indented. Just underneath, Figure 2E., is a free-hand drawing of an interphase chromosome (interphase S) region, with dimensions, where these variations would result in rough, not smooth, overall surface. Figure 2F&G. shows that the prophase chromosome structure (prophase S’) also had variations, primarily variations in the size of the pleats/indentations, which creates wider or more narrow X axis structure. Figure 2H & I. show that the mitotic chromosome coiling (mitotic S’’ using the additional coiling) would have mitotic diameter variations because different sized prophase variants were used to coil. The left side coiling used the prophase chromosome dimensioned from the left side of Figure 2F, while the right side used the prophase chromosome of Figure 2G. Note, while the mitotic diameter varies, the inside diameter is constant.

### A Proposal for Interphase S, Prophase S, and Mitotic S’’ as Unified Chromosomes Structures Throughout the Cell Cycle

The fundamental architecture of chromosomes must allow, in a seamless fashion, the dramatic changes in their overall structure to take place during the cell cycle: condensation in prophase to form the mitotic chromosome structure and decondensation in telophase as the interphase chromosomes/ nucleus reforms. As a first approach to understanding this change in chromosome structure, we modeled human chromosome 21, the smallest human autosome (45 million base pairs long with 196,000 nucleosomes, Table 1), as a plausible architectural route through the cell cycle (Figure. 3). Starting in interphase (G1) for this specific human chromosome, chromosome 21, the interphase S also folds into 14 micron sized large-scale coils (Figure. 3A& legend). We conjecture that all interphase S structures are associated with large-scale loops, plausibly involving mega-loops as a fundamental aspect of their structure. Here we define such loops as “Primordial Coils”. Prophase S’ shows prophase chromosomes, now condensed as described (22), but with the same number and spacing as the Primordial Coils in interphase S (Figure 3B). Highlighting the assembly of mitotic chromosomes, by visualization (Figure. 3C movie), readily reveals the dynamics of mitotic chromosomes coiling. Specifically, we conjecture that mitotic coiling created two tighter mitotic coils (about 500nm in diameter) for every Primordial Coil, for a total of 28 mitotic coils. Played backwards, the movie highlights telophase. We propose that in telophase, mitotic chromosomes uncoil discontinuously. Some spaced coils remain--possibly two adjacent mitotic coils fused to create micron size Primordial Coils. Such Primordial Coils remain in interphase/G1 to instruct the assembly of chromosome territories (Figure 4A-C &D). Thus, these observations indicate that special coils—Primordial Coils--build features that architecturally are ready to proceed to the next step in the cell cycle.

**Figure 3.**
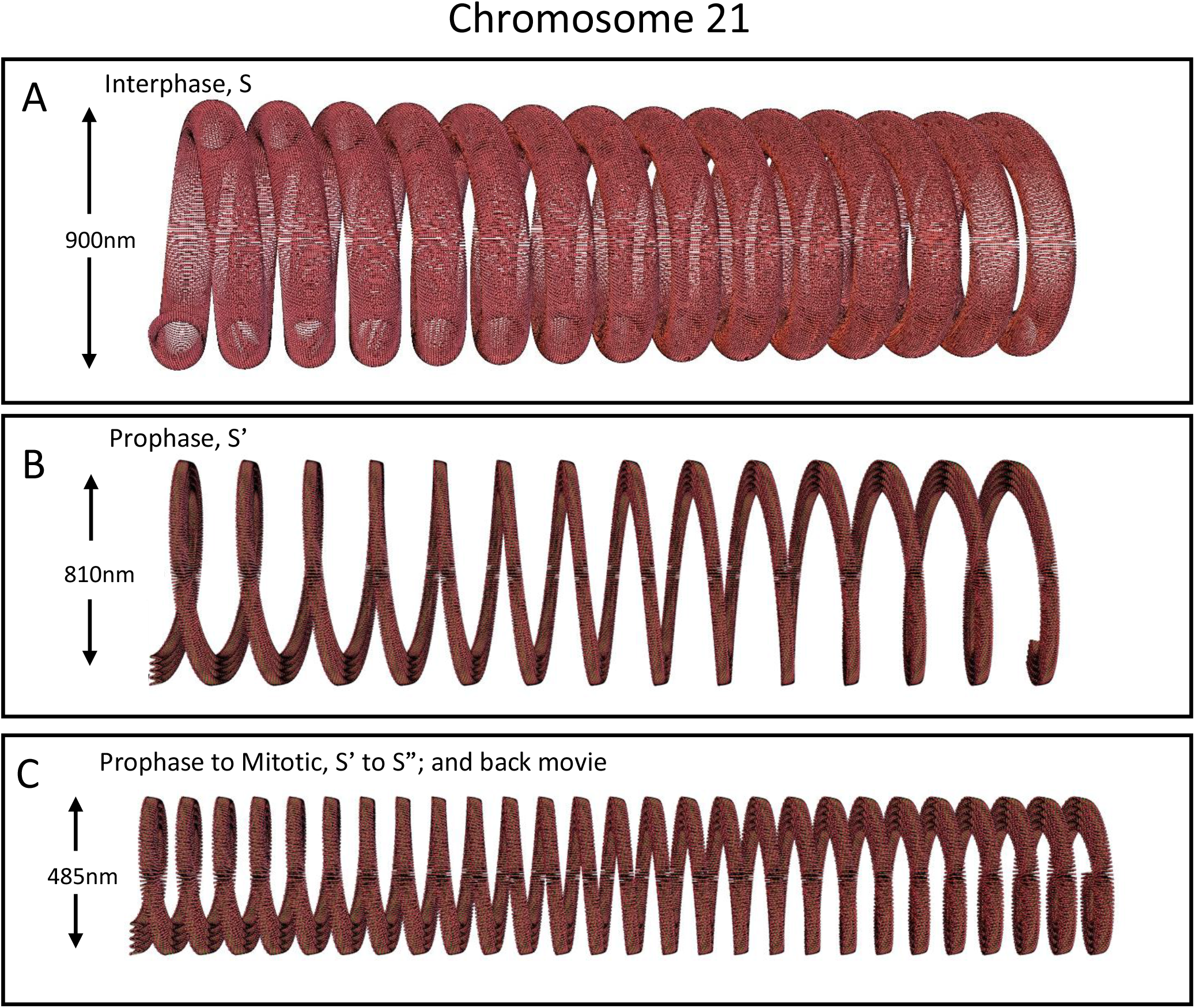
A unified hierarchical coiled chromosome architecture transits through the cell cycle, as depicted for human chromosome 21. Panel A depicts chromosome 21 as an interphase structure with 14 approximately 1micron large coiled loops. These coils resulted from a partial specific uncoiling of the mitotic chromosome in telophase (see panel C), and are called Primordial Coils to emphasize their organizational relationship in the cell cycle; these coils were the basis of chromosome territories described in Figure 4. Below the level of the Primordial Coils is the interphase chromosome structure, helically coiled structure of the nucleosome 11nm fiber, interphase S, extensively described in (20) and Figure 1. Panel B depicts chromosome 21 as a prophase chromosome structure, a compressed and pleated structure (prophase S’), described in (22) and Figure 1. In prophase, this prophase chromosome has the same locations and numbers of Primordial Coils (14, 1micron) resulting from the telophase mitotic chromosome unwinding. Panel C is a movie (clickable of the structure as described in Figure 1) depicting the further compaction of chromosome 21 from a prophase structure (the first frames of the movie) coiling into a chromosome 21 mitotic chromosome--forming the tighter 0.5micron mitotic coils. The mitotic chromosome architecture was described in (22) and Figure 1. It can be seen that two mitotic coils form for each Primordial Coil leading to 28 final mitotic coils. It can be pointed out that the mitotic chromosome for chromosome 21, seen here, is slightly extended to emphasize the coiling, if compressed a correct sized chromosome 21 would result (22). Panel C also shows the telophase chromosome structure as part of this phase of the cell cycle. The movie is just played in reverse; the movie bar is now moved right-to-left and chromosome 21 unwinds specifically two mitotic coils for every Primordial Coils, recapitulating what one sees for prophase and interphase. Initially, this unwinding of the mitotic chromosome is a prophase structure, which rapidly turns into an interphase structure. Going from Figure 3C to Figure 3A suggests that the cell cycle has nested coiling— architectural--features that allow the chromosome to process through the cell cycle seamlessly.

**Figure 4.**
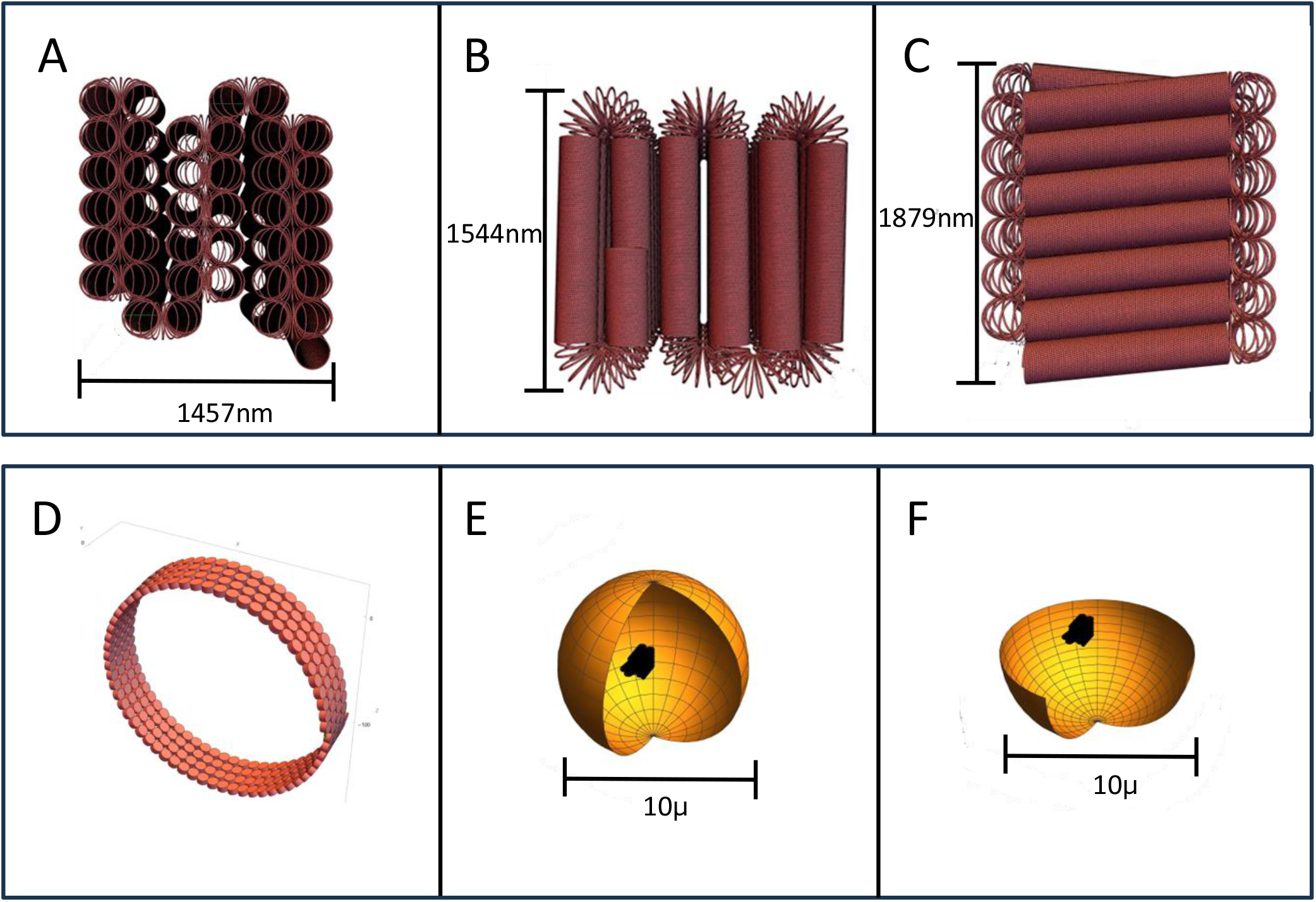
The helical coiling architecture is used to construct higher eukaryote micron sized Chromosome territories. Figure 4A-C. depicts human chromosome 19 constructed as human chromosome territory, in X,Y,&Z views with dimensions. A chromosome territory is built by using its interphase chromosome 1micron loose Primordial Coils left over from specific unwinding, in telophase, of the mitotic chromosome. These loose micron sized coils are flattened to reduce space with further compaction to make chromosome territories. The micron loops are further defined in Figure 3. however, they show that the residual telophase mitotic chromosome unwinding is used architecturally for chromosome territories. Figure 4D. shows a movie (click on the center of interphase S loops) of the construction/organization of chromosome 19 chromosome territory. Figure 4E &F. depicts chromosome 21, as another example chromosome territory, as one homologue, correctly dimensioned, in a 10micron (average size) nucleus diameter.

### Interphase S Chromosomes as Chromosome Territories

Interphase chromosomes are not intertwined randomly in the nucleus but occupy distinct regions known as chromosome territories (38-41). The chromosomes in most organisms that have been examined are arranged in the Rabl configuration (named after the 19th-century cytologist Carl Rabl), in which the centrosomes cluster at one side of the nuclear envelope and the telomeres are attached to the opposite side, a consequence of chromosome segregation at anaphase (42-47). . We recently built an interphase S chromosome as a Rabl chromosome territory (20).

Higher eukaryotic genomes are segregated into many individual chromosomes, and organize their chromosome territories as micron sized patches (39). Initially, we suggest a framework for chromosome territories. This framework is a series of approximately 1micron Primordial Coils (Figure 3). These Primordial Coils, are straightened then tightened, to make an approximately micron sized patch (Figure 4A-C). Human chromosome 19 is the representative example for the structure of the patch sized chromosome territory (Figure. 4A-C). Visualization reveals possible kinetics that underpins the assembly of chromosome 19 into a chromosome territory (Figure. 4D movie). We also modeled the chromosome territory of human chromosome 21 and show how it fits into a correctly scaled 10 µm diameter nucleus (Figure 4E and F)

All 46 human chromosomes, as patch chromosome territories, have to fit into a 10micron (average size) nucleus. Using chromosome 19 as a volume template for a chromosome territory, and the 46 chromosomes (calculated as nucleosomes/chromosome) allows possible scaling of the chromosome territories (Table 1). The chromosome territories sizes, located in Table1,column 5, are summed together (times two for the other homologues); it was possible to show that all 46 chromosomes fill a 10micron nucleus at the 80.29% volume level. This is likely an upper limit as we assumed cubic chromosome territory volumes (see Figure 4A-C.). This result suggests that interphase S architecture is compatible with patch-based chromosome territories. Given the remaining 20% (or more) volume in the 10micron nucleus, we expect that there will be freedom to uncoil/distribute the chromosome territories in more open configurations, for example to enable interphase S for transcription. Chromosome motion should expand /unfold somewhat the chromosome territories. We emphasize that the patch chromosome territory, as built here, is a model for that assembly of chromosome territories, but that other chromosome territory models are plausible.

The individual chromosome territories in our model are quite dense. The chromosome 19, for example, contains 268,000 nucleosomes (Table 1) in a patch that spans approximately 4 µm^3^. Such a configuration should permit assembly of topologically associating domains(TADs), as regions are close together. Indeed, in our STEM Cryo-EM Tomography images we noticed several side-by-side parallel associations of interphase S regions (20). Moreover, live cell imaging studies revealed predominantly intrachromosomal loop associations rather than interchromosomal interactions (42).

### Implications for DNA Replication Involving Interphase S, Prophase S’ and Mitotic S’’ Chromosomes

Any chromosome architecture has to take-into-account the impact of DNA replication (48-51). We show that after replication the two interphase S structures interdigitate-nest-inside each other, a natural feature for Slinky architecture (Figure 5A). Upon replication completion, the two interphase S structures could slide out from each other, likely still positioned within close spatial proximity from one another. Another important feature of the nested interphase S Slinky structure is that they are able to rotate about the long axis of the helix allowing structural flexibility. We note that it is conceivable that strand separation occurs at the start of G2/prophase or alternatively that it is discontinuous because of replication timing. A sight complication is seen at centromeres (52). We hypothesize that the replicated centromere DNA interphase S structures still interdigitate/nesting held together by cohesin (53-56) (Figure 5B). It is plausible that late DNA replication of the centromere may play a role in this process.

**Figure 5.**
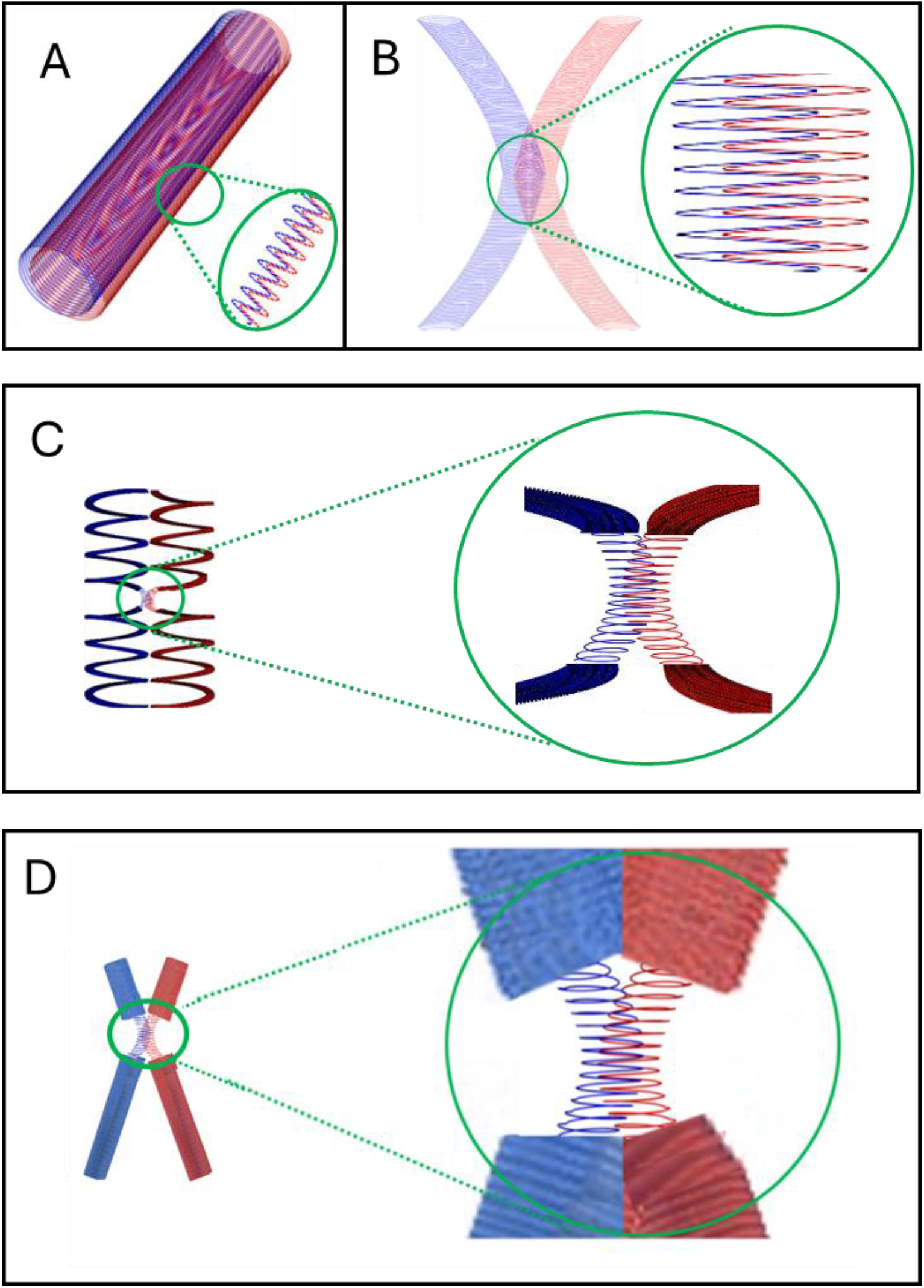
Helical coiled architecture as a function of chromosome replication. Panel A depicts, as line drawing, two interphase (interphase S) replicated chromosomes, blue line for the original, and red for the replicated interphase chromosome; just after replication the two chromosomes are nested, a natural Slinky helix process--inside one another as the insert shows. The insert is an enlarged view of the nesting. After replication, the two replicated helices just slide out from each other, seamlessly, as seen in Figure 5B. Figure 5B shows that the centromeres are treated differently; they continue to interdigitated because they are held together by cohesin protein (53-56). Figure 5C. shows prophase replicated chromosomes, blue for original and red for replicated, fully separated except for the centromere. These still are held together by cohesin and interdigitated. Figure 5D. shows replicated mitotic chromosomes still held together by the cohesin mediated interdigitation until anaphase where the cohesin enzymatically releases (75-78). These are accurate human chromosome 10 (23), except for the centromere regions.

The separated replicated interphase S chromosomes are free to form prophase chromosomes, as prophase S’, at the right time in the cell cycle, as shown in Figure 5C. Again, the slight complication is the centromere region; centromeres remain likely uncondensed as interphase S, interdigitated/nested together, held tightly by cohesin protein (53-56), and diagrammed in Figure 5C insert.

Mitotic chromosomes coil as part of the mitotic S’’ coiling, with the replicated sister chromosomes side by side shown in Figure 5D. Also, there is the centromere complication; the two sister centromeres are still in interphase S, coiled, interdigitated/nested together (Figure 5D insert). The key point is that the centromere, in interphase S, is not further coiled to prophase S’ or mitotic S”, and the sister chromosomes, held together by cohesin protein, associate as a pair as seen in the chromosome spreads (26-29). Mitosis takes place with the attached mitotic chromosomes, a lengthy process (49,57), until at anaphase cohesin is cleaved to release the two sister chromosomes (53-56).

## Discussion

### Coiling Aspect

Coiling helically, in some cases sequentially, seems to be a paradigm that is used over and over again to describe folding patterns for DNA structures. For example, DNA folds as a right-handed two-strand-helically coiled structure. The nucleosome is organized as a discontinuous left-handed coiled structure that is folded around the octamer histone core (58). Helically coiling involving the nucleosome 11nm fiber, in turn, assembles into an interphase S structure. A proposal, in a previous paper, showed that polytene chromosomes, a representative--with higher order structure, -- interphase chromosome(G1/S), could be built with coiling features of the interphase S structure(discussed in 20). Prophase S’ is modified interphase S, still with its coiling, and this could be extended to lampbrush prophase I chromosome structure as an example(20). Helically coiling prophase S’ to make mitotic S’’ (hand unspecified) along the same line follows the coiling paradigm. How is it possible to coil this plethora of structures, and which enzymes/proteins are required? These are important study directions.

We recently showed that condensin, which functions in chromosome condensation and segregation in mitosis and meiosis (59,60), with just the right molecular spacing, was possibly involved in the assembly of prophase S’ topologies (20). Prophase S’, as built and displayed, involved a two-step process, because the software for the correct molecular and structure simulations was somewhat inflexible. Prophase S’ could be envisioned, in vivo, as a much simpler, possibly enzymatic/protein problem. In interphase S, on each side of the interphase S gyri at eight separate spaced points--four points at the top and four points at the bottom of the gyrus—could be identified then pulled using possibly concatenated condensin to both collapse the interphase S gyrus into a race-track configuration associated with deep indentations (see Figure1B) (59,60).

### Interphase S Polymer Considerations

We emphasize that interphase S, the interphase chromosome architecture, is in essence a special polymer structure that is folded into an approximately 200nm diameter coiled, hollow, and flexible Slinky. It is likely to follow, polymer statistics (1-6&61,62), though special attributes apply, especially since this is a densely coiled structure with special features. For example, in one axis interphase S is very easily bent and deformed, while in the other directions the interphase S gyri compress restricting bending. How the polymer dynamics are affected is an interesting question, and complicated by the very high density of nucleosomes, in some cases akin to that of chromosome territories chromosomes. The issue of a somewhat small 10micron nucleus, constraining the chromosomes, which has to effect polymer statistics. In summary, the interphase S architecture, overall, has many features in common to the polymer-based systems, physics, and thinking the community and literature currently considers (1-6&61,62).

There are several attributes to consider for interphase S structures. First, interphase S are dense coils that are predominantly perpendicular to the interphase S long axis and are very densely packed (approximately 64 nucleosomes—12.8kb/gyrus). Second, interphase S are hollow, and when pulled out almost free diffusion of the hollow interior compared to the interphase S exterior; If interphase S is compressed as it is for heterochromatin, the hollow interphase S interior would have restricted diffusion, possibly used for protein storage or a different enzymology. Third, interphase S structures can be straight for regions, and the closely packed interphase S gyri could associate one gyrus with another giving rise, possibly to better rigidity (20). Fourth we note that the exterior surface of interphase S is extended compared to the hollow interior and long range 2-dimensional aligned phased nucleosome interactions are possible (20,22). It is conceivable that this would enable increased cooperative interactions involving chromosomal-associating proteins (63,64). Fifth, interphase S allows for adjacent interphase S coil nesting interactions as pointed out in the interphase S replication and Centromere adhesion situations. TAD interactions could make use of such an interphase S feature. Sixth, interphase S structures could well allow for genetic coordination by epigenetic marks on adjacent interphase S gyri. Seventh, it is conceivable that such a delicate interphase S structure is potentially sensitive to fixation making it difficult to interpret structural information derived, for example, from formaldehyde-fixed cells.

### Structure Determination Aspects

A major point to make is that additional experimental data is urgently needed. The detailed structures for prophase S’ or mitotic S’’ are not existent. The right experimental system, we feel, is cryo-EM Tomography, since the structures are likely faithfully preserved in a glassy frozen aqueous state. Recent progress in STEM EM technology will allow even better data collection and computer processing; double tilts, on orthogonal axis, followed by deconvolution shows improved Z resolution; better filling of the missing wedge coming from incomplete high angle tilts information (65). Recent detector technology now provides fast 96x96 pixels at 120,000 frames per sec. data collection, possibly allowing increased contrast or possible specific atom detection (like Nitrogen for nucleic acid/DNA, for example 65).

Detailed 3-Dimensional optical microscopy will also greatly help prophase S’ and mitotic S’’ structure determination. In this regard, the polarization microscopy/birefringence optical studies of nuclei, suggesting ordered structures, may need to be re-examined; the nuclear substructure, never interpreted or understood (66), is reminiscent of the interphase S architecture described in (20)

### Variability in Interphase S, Prophase S’ and Mitotic S’’ Structures

Since it is possible to vary each of the interphase S, prophase S’ and mitotic S’’ structures, one could ask why does such a degree of variation exist? Is this variation used to distinguish specific genetic regions from one another involving differences in transcription timing or regulation? One hypothesis comes from observations made in Drosophila melanogaster. All 5000 polytene bands are structured into distinctive shapes and sizes (67). Recent studies showed that polytene bands are folded as TADs ( the same as diploid interphase chromosomes ), and are comprised of tissue distinctive regulatory elements (68,69). Hence, we conjecture that interphase S variations consist of DNA elements with distinct regulatory functions (71). We again emphasize that polytene chromosomes are examples of bonified interphase chromosome structure(67-70).

### Interphase S Replication Aspects

DNA replication process suggests that the interphase S architecture facilitates replication segregation and possible control of the replication events (see Figure 3). The centromere on mitotic chromosomes, for example on metacentric chromosomes, is a constriction, compatible with interphase S structure. The centromere adhesion for prophase and mitotic chromosomes was an old problem, with a suggested molecular solution.

### Homologue Chromosome Considerations

An important experimental result showing restricted nuclear chromosome localization needs discussion. Hua and Mikawa documented that in low passage human primary cells (but not cancer cell lines) the two homologues are restricted to separate halves of the nucleus, not intermixed, at least during early metaphase and anaphase (71). A recent study provide evidence that chromosome centromere markers components, are separated by a deep cleft in the middle of the metaphase chromosome mass along the centrosome axis of the nucleus, suggesting restriction of homologues in significant phases of the cell cycle, with little or no mixing of homologous chromosomes (72).

### Conclusions

In conclusion, we note that a Slinky based architecture is able to withstand potential complications—kinetics, structure variants, flow through the cell cycle, micron patch chromosome territory dimensions and chromosome replication issues, indicative of architectural robustness. However, insights into mechanisms that permit the folding of 2m of DNA, packed with nucleosomes, into a 10micron mammalian nucleus and proceed orderly through the cell cycle remain rudimentary and will require further experimentation using both cryo-EM approaches as well as live cell imaging.

## Methods and Materials

Computer modeling software, and computer hardware/software utilized for this paper, follows our previous publications (20,22)

The computer modeling of the chromosomes, at all levels, utilized a software package written by an engineering group (30) that allows sequential helical coiling of defined sized structures. This package is run under, and requires, Mathematica 12.3 (73) under Linux in large work stations. Mathematica 12.3 was run under Windows 7, Intel Core i5 CPU 3.20 GHz processor with 4 cores and displayed on a Samsung C27F591monitor and NVIDIA Quadro K1200 video card and 8 GB of memory. Mathematica was also run on a CentOS 7.3 Linux system running on Intel Core i7 CPU 2.80 GHz processor with 16 GB of memory. The display of the Linux generated output was visualized on the Windows 7 computer.

The software was modified to take-into-account the time dependent/kinetic interphase S configuration, prophase S’ modifications, and the mitotic S’’ coiling. The movies of the chromosome timeline were all made with Wolfram Mathematica 13.0. Each frame of the movie was generated as a Mathematica Graphics3D figure and stored in an array. The parameters of the Graphics3D command were changed in each frame to reflect the conformational changes of the chromatin structure. Finally, the array of stored frames is converted and stored as to a QuickTime movie using the Mathematica Export command.

The various scripts for the software are supplied by the authors upon request.

## Acknowledgments

We gratefully acknowledge the advice and suggestions of Professors Thomas Cremer, Geeta Narlikar, Zvi Kam, Lisa Hua, Lloyd Smith, and Marc Shuman. We are grateful for the editing of Carol Featherstone of Life Science Editors. The computing research and computer resources are self-funded by A.M. and J.S. Research in the Murre laboratory is supported by the National Institutes of Health RO1AI082859, RO1AI082850 and US1-BSF 2019280.

